# Parapipe: a Pipeline for Handling Parasite NGS Datasets and its Application to Cryptosporidium

**DOI:** 10.1101/2025.01.16.633364

**Authors:** Arthur V. Morris, Guy Robinson, Rachel Chalmers, Simone Caccio, Tom Connor

## Abstract

*Cryptosporidium*, a protozoan parasite of significant public health concern, is responsible for severe diarrheal diseases, particularly in immunocompromised individuals and young children in resource-limited settings. Analysis of whole genome next generation sequencing (NGS) data is a critical next step in improving our understanding of *Cryptosporidium* epidemiology, transmission dynamics, and genetic diversity. However, effective analysis of NGS data in a public health context necessitates the development of robust, validated bioinformatics tools. Here, we present Parapipe, a modular ISO accreditable bioinformatics pipeline designed for high-throughput processing and analysis of *Cryptosporidium* NGS datasets. Built using Nextflow DSL2 and containerized with Singularity, Parapipe is portable, scalable, and capable of end-to-end analyses, including quality control, variant calling, multiplicity of infection (MOI) investigations and phylogenomic clustering analysis.

Using both simulated and real-world datasets, we demonstrate Parapipe’s ability to resolve genetic heterogeneity, identify mixed infections, and generate high-resolution phylogenomic insights. Here, we use it to carry out a comparison between whole genome single nucleotide polymorphism (wgSNP) typing and the conventionally used gp60 molecular typing scheme. Compared to existing pipelines, Parapipe uniquely integrates MOI analysis, enabling the differentiation of mixed infections and supporting epidemiological investigations. Parapipe’s design facilitates integration with geographic, demographic, epidemiological and environmental data, enhancing its utility for tracking transmission pathways and outbreak sources.

Parapipe represents a significant advance in utilising genomics for public health surveillance of *Cryptosporidium*, offering a streamlined and reproducible framework for analysis with potential application to other pathogenic protozoa. By automating complex workflows and enabling detailed genomic characterization, Parapipe provides a valuable tool for public health agencies and researchers, supporting efforts to mitigate the global burden of cryptosporidiosis.

## 1.0 Introduction

*Cryptosporidium* is a single celled Apicomplexan parasite genus that is a leading cause of diarrhoeal disease in humans and animals. Over 200,000 deaths of children annually in Asia and Sub-Saharan Africa alone are attributable to this parasite [1]. Young, malnourished, or otherwise immunocompromised individuals are at particular risk of significant morbidity and mortality [2]. Humans acquire the infection faeco-orally through multiple routes and vehicles, including direct human-to-human and animal-to-human contacts and consumption of contaminated water and food.

Although at least 48 species of *Cryptosporidium* have been described, infection in humans is mostly caused by two species: the zoonotic *C. parvum* and the anthroponotic *C. hominis*. Currently, there are no truly effective drugs or vaccines to treat or prevent cryptosporidiosis, and thus control is heavily dependent on the prevention of infection, which in turn requires a detailed understanding of Cryptosporidium epidemiology, population structure, and transmission dynamics [2]. In this context, genomic analysis could provide a cornerstone for the control and prevention of cryptosporidiosis, as it allows for high-resolution typing evaluation, beyond that which is possible using conventional molecular techniques. However, a pre-requisite for the routine use of genomics for human and animal public health purposes necessitates the development of reliable bioinformatic pipelines, validated to a high standard, which enables data analysis to be completed in a short space of time. The timely development of an appropriate pipeline coincides with technologies enabling next generation sequencing (NGS) of *Cryptosporidium* from DNA extracted directly from stools using hybridisation baits [3], [4]. Previously, only limited numbers of specifically selected specimens have been genome sequenced from the oocyst (transmissive) stage of this practically non-culturable organism that is present in low density in complex matrices such as faeces [5]. *C. parvum* is conventionally subtyped by interrogating fragment length and sequence variation across a highly polymorphic Variable Number Tandem Repeat (VNTR) region within a surface glycoprotein gene, gp60[6]. This locus has been utilised for Cryptosporidium subtyping for over two decades and provides the only standardised subtyping scheme for *Cryptosporidium* [You’ll need to find a NEW REF 8]. However, despite limitations of gp60 subtyping (single locus, usually by Sanger sequencing), its undeniable utility, comparatively low cost, wet lab simplicity, and the familiarity of this subtyping scheme within the wider Cryptosporidium community, has encouraged its continuing use [5], [7]. Yet there is evidence that higher discrimination, improved epidemiological concordance, identification of multiplicity of infection, and greater public health utility can be provided by multilocus genotyping [8], [9].

Here, we present Parapipe, a bioinformatic pipeline that has been developed to a standard that would allow it to be accredited as part of an ISO 15189 or 17025 laboratory process, and meet the standards and requirements of public health microbiological investigations [2]. Parapipe has been developed for *Cryptosporidium* to meet the needs of clinical laboratories, and is intended to solve the issues of NGS data quality assessment and control, mapping, variant calling, multiplicity-of-infection investigation, and clustering analysis. Following the bioinformatics approach adopted for other accredited genomics services within Public Health Wales, we have developed a modular system, in which Parapipe is subdivided into functional components that can be tested individually, as well as part of the larger whole. We carry out extensive module and end-to-end testing to demonstrate pipeline reliability and use Parapipe to investigate the heterogeneity and relatedness of a *C. parvum* NGS dataset.

Parapipe represents the first publicly available pipeline designed to carry out essential preprocessing and analysis of *Cryptosporidium* NGS data. Although Parapipe was designed with *Cryptosporidium* in mind, in principle, it can also be used on a wide range of pathogenic eukaryote and prokaryote taxa.

## 2.0 Materials and Methods

### 2.1 Implementation

Parapipe is a command line-based pipeline written in Nextflow DSL2 [10], allowing for modularisation and containerisation using Singularity [11]. The use of Nextflow and Singularity ensures that the software is portable between systems, and enables a simplified process for installing and running the pipeline, either using on-premises systems or on a commercial cloud. Parapipe is built using well-established read pre-processing, quality control, mapping, and variant calling tools. Parapipe is a linear pipeline, whereby each module is executed in a prescribed order on every sample.

As input data, it takes a set of paired end reads in FASTQ format. Each read set is passed through the pipeline as an individual run, allowing multiple read sets to be processed in a parallel manner. Reference datasets (consisting of a sequence and annotation pair) can be deposited in a subdirectory within the Parapipe root directory, for convenient use by the user via a command line argument.

Parapipe consists of two primary work modules, comprising twelve individual processes (1.1 to 1.8 and 2.1 to 2.4, see Figure 1). The workflow from module 1 to module 2 is automated. The first module prepares the reference data, and performs quality control, pre-processing, and mapping on the input reads. The second module performs variant calling, clustering, phylogenetic, and phylogenomic analysis.

**Figure 1.**
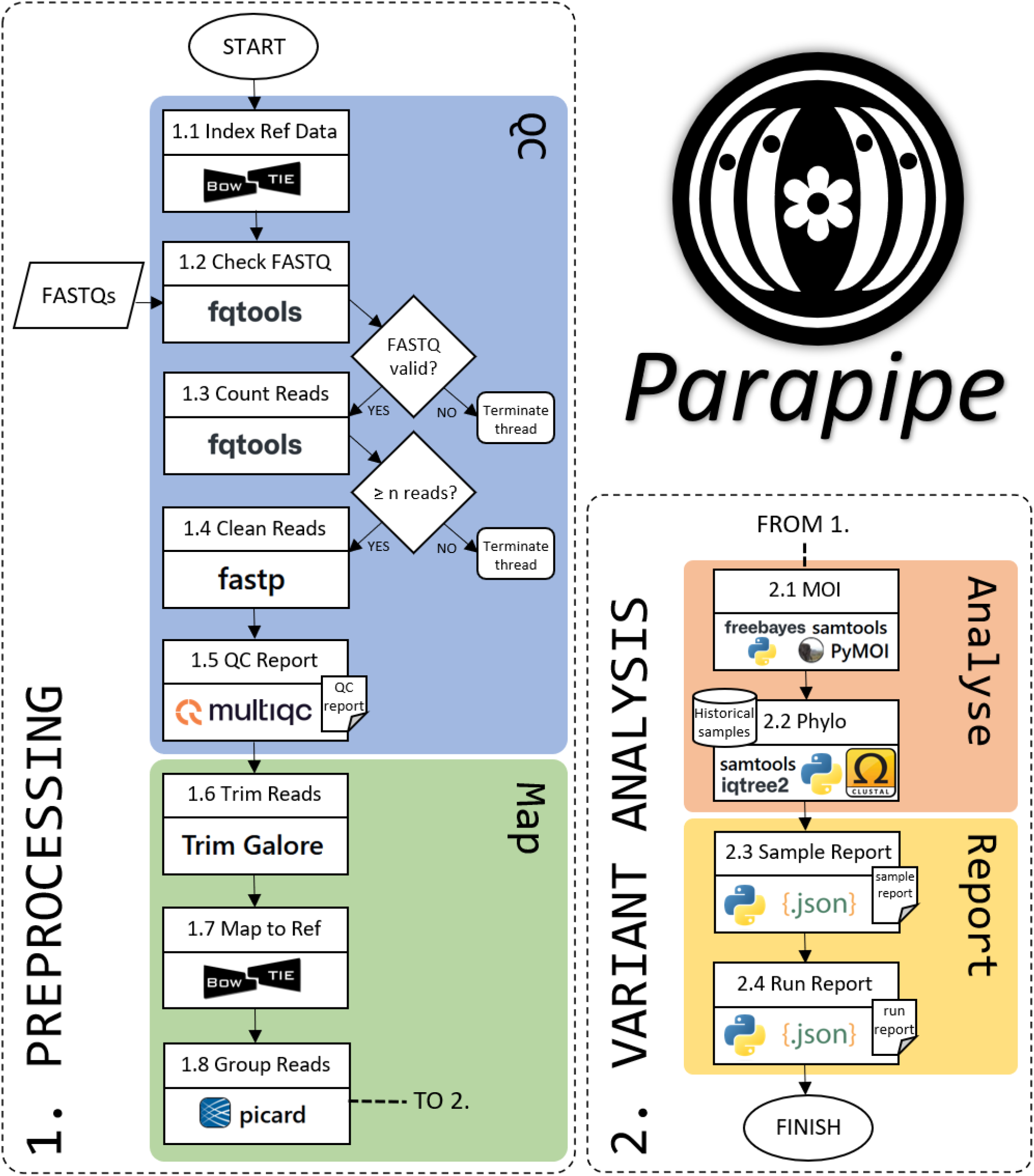
Workflow schematic for Parapipe, detailing the processes that comprise the two primary work modules within the pipeline, preprocessing and variant analysis.

#### 2.1.1 Module 1

Module 1 consists of eight processes with two logic gates. Process 1.1 fetches and prepares the reference files by constructing a `Bowtie2` index and a `samtools` faidx index from the reference FASTA file [12], [13]. FASTQ files are then checked using `fqtools` (process 1.2 and 1.3) to ensure the input FASTQ files are valid and contain enough reads (by default, this threshold is 1 million reads) [14]. Logic gates exist to evaluate the output of `fqtools`. FASTQ file sets that do not pass these checks are not processed further, and the relevant threads are terminated. Processes 1.4-1.5 perform cleaning and quality control of the input FASTQ sets using `fastp`and `fastQC` and aggregates all QC reports from all FASTQ sets in this Parapipe run using `multiQC` [15], [16]. Read trimming and mapping using `trimGalore` and `Bowtie2` respectively are carried out during processes 1.6 and 1.7 [17]. The BAM files produced by process 1.7 are deduplicated and assigned a group using `Picard` (process 1.8) [18].Since the input read sets constitute a single run of an isolate, all reads in each sample FASTQ set are assigned to the same read group. This is a necessary step to facilitate downstream variant and sample heterogeneity analysis.

#### 2.1.2 Module 2

Module 2 carries out the variant analysis and comparative genomics within Parapipe. Process carries out Multiplicity of Infection (MOI) investigation to elucidate the extent to which multiple distinct genomic variants exist within the dataset. MOI can occur as a result of either a mixed infection, or as a product of individual oocysts containing genomically heterogeneous sporozoites [5]. This process is driven by the `PyMOI` library for Python3 and the R library `moimix` which carry out haplotype analysis using the full SNP repertoire [19]. Datasets are subjected to variant calling to identify SNPs using `FreeBayes` (process 2.1) [20]. The full repertoire of wgSNPs across each sample is then used to construct a tree and Principal Co-ordinate Analysis (PCoA) clustering plots representing distance in SNP space (process 2.2).

SNP space plots are generated using Python libraries (`scipy` and `matplotlib`) [21], [22]. Finally, reports pooling relevant data for the entire run (process 2.4), along with more granular reports for each sample (process 2.3) are constructed. These reports contain a table of mapping statistics, total and unique SNP count for each sample, data produced from the MOI analysis (process 2.1), and phylogenetic analysis (process 2.2).

All relevant data are deposited in an output directory, the structure for which can be found in the supplementary materials section.

#### 2.1.3 Reporting

Two kinds of report are produced by Parapipe. First, a report is produced for each sample, containing a table detailing the results of quality control, mapping, MOI analysis, and SNP analysis, along with phylogenetic plots for SNP distance. A further report is produced which pools all relevant data for the entire run. The sample reports are provided in PDF format. The run report is provided in both PDF and HTML format. JSON files, which are machine readable, for each report are also provided to the user in the output run directory.

### 2.2 Evaluation

#### 2.2.1 Read Simulation

All reads were simulated using ART v2.5.8 [23] using the Illumina HiSeq 25 (HS25) profile with read length of 150 bp and insert size of 200 bp. Reads were simulated as pairs with the average quality score of all reads set to 30. Reads were simulated using the updated *Cryptosporidium parvum* Iowa II-ATCC reference genome (GenBank assembly GCA_015245375.1) [24].

#### 2.2.2 Module Testing

Module testing was carried out using simulated raw read data to test each of the functional components of Parapipe, outlined in Fig. 1, to ensure reliability. The *C. parvum* reference genome (Iowa II-ATCC) [24] was used as a base sequence into which variants can be introduced and reads simulated to test whether Parapipe can reconstruct known and controlled phylogenetic relationships and heterogeneity profiles. Processes within Parapipe were grouped into testing modules (TMs) according to whether they carry out a single testable function within the pipeline. Each TM within the pipeline was tested using a set of simulated reads designed to test the functional element of that module. Further details of the testing modules can be found in the supplementary material (see supplementary Tables S1 and S2). For example, TM05 refers to the module that investigates sample heterogeneity, using the python library `PyMOI` and the R package `moimix`(process 1.8-2.1). The test dataset was simulated using the *C. parvum* reference genome (IowaII ATCC) as a backbone to construct a read set covering the genome to 10x in BAM format, and either 100 (representing low level of heterogeneity) or 1000 (representing a moderate level of heterogeneity) SNPs introduced at equally distributed random locations across the genome to meet specified allele frequencies.

Table S2 details the simulated dataset built to test the capacity of Parapipe to compare input samples by the presence of SNPs. This dataset was designed in a hierarchical manner: there are two lineage orders, such that each nth order lineage (e.g. L1.2 would be a second order lineage) exhibits all SNPs of <nth ordered lineages on the same branch (e.g., L1.2 would exhibit all the same SNPs as L1, with a further set of L1.2 specific SNPs). The module used to carry out phylogenomic inference in Parapipe makes no such assumptions about hierarchy or SNP inheritance, since this could result in fallacious evolutionary relationship inference (i.e., it does not necessarily follow that a population with a set of SNPs is the ancestral population of one which also exhibits them in addition to a novel set of SNPs).

Each simulated dataset was run through only the module it was intended to test, with prior processes that may alter or affect the results removed to ensure validity of results. Only processes producing output required to run the test module were executed.

Results from Module Testing can be found in the supplementary materials.

#### 2.2.3 End-to-End Testing

Testing of the full pipeline as a single unit was carried out using a *C. parvum* dataset consisting of simulated raw read data of both mixed and clonal samples (n=22). The dataset consists of three discrete lineages down to 2nd order, each consisting of a single dataset representing the first order lineage, and five datasets representing 2nd order lineages (n=18). Compound datasets were also constructed, representing a number of first order lineage mixtures (n=4). Datasets were simulated using a depth of 10x.

#### 2.2.4 Testing on Real Data

Parapipe was run on a dataset consisting of 20 *C. parvum* whole genome samples derived from human and animal infections from four European countries belonging to five gp60 subtypes from gp60 family IIa. Phylogenetic trees were generated using the wgSNP repertoire, and the gp60 sequence. A description of the dataset can be found in the supplementary materials. A minimum read number threshold of 100,000 was utilised during quality control. The minimum fraction of the reference genome that must be covered for inclusion in phylogenetic analysis was 80% to a depth of at least 5x. For MOI investigation only alleles with a minimum frequency of 0.05 and covered to a depth of at least 5x were used.

#### 2.2.5 Run Time

Parapipe processed the 22 samples representing the end-to-end testing dataset (32838480 reads total) in 1 h 9 min 11 s (9.2 CPU hours), and a single dataset (1216240 reads total) in 5 m 3 s (13 CPU min). All testing was carried out on a laptop running Ubuntu v20.04.6 with 11 cores and 30Gb physical memory, and using Nextflow version 24.10.1.

#### 2.2.6 Comparison Against Other Pipelines

The majority of the pipelines designed to process and analyse pathogen NGS data are specific to viral and bacterial genomes. Due to the domain specific challenges and needs faced by protozoan genomics, those pipelines are not suitable for processing *Cryptosporidium* NGS data. Nonetheless, the pre-assembly functionality of these pipelines was compared to that of Parapipe.

## 3.0 Results

Running Parapipe on the end-to-end test dataset recovered both the 3 major lineages (L1, L2, and L3), and the hybrid lineages simulated within this dataset. Results of whole genome SNP-space analysis showed three major clusters, along with four satellite points (L1∪2, L1∪3, L2∪3, L1∪2∪3).

Inspection of the PCoA plot (Figure 2) revealed more detailed clustering information. Lineages 1, 2, and 3 all form discrete clusters within the dataset. L1∪2 presents as equidistant between lineage clusters 1 and 2, L2∪3 lies between lineage clusters 2 and 3, and L1∪3 lies between lineage clusters 1 and 3. Furthermore, L1∪2∪3 lands directly in the middle of lineage clusters 1, 2 and 3. All satellite samples show clear indication of sample heterogeneity (*Fws*>=0.95), with L1∪2, L2∪3, and L1∪3 showing a single allele frequency band centring at 0.5, indicating the presence of 2 distinct populations (each exhibiting an *Fws*>=1.0). Dataset L1∪2∪3 presented with two bands centering at 0.5 and 0.75, indicating the presence of more than two populations (*Fws*>=1.0). All samples covered >99.9% of the reference genome with a median DOC of between 20-59x, with high equality of read distribution (Normalised GG area of 0.935-0.96).

**Figure 2.**
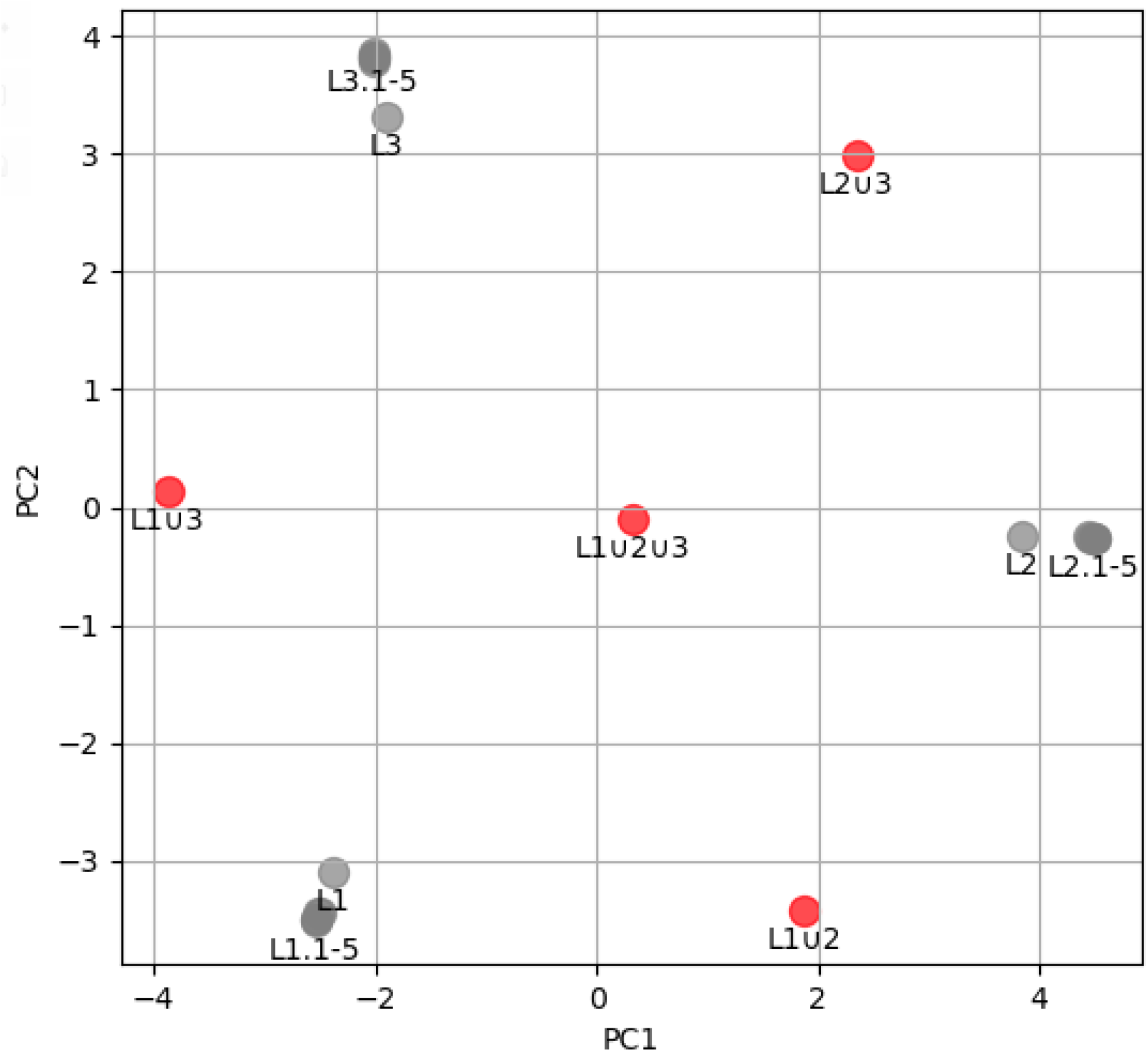
A PCoA plot of the wgSNP distance generated from the end-to-end dataset. PC1 plotted against PC2 with 74% of the variation space captured across these two components. Marker legend: Red=mixed, grey=clonal.

Using Parapipe revealed substantial population structure within the *C. parvum* IIa dataset. Analysis by wgSNP Hamming distance indicated the presence of two genetically diverse groups (C1 and C2), which showed association by geographical origin (Fig. 4, panel A). Group C2 could be further subdivided into 3 subgroups (C2a-c). Diversity within these groups was considerable and showed some correlation within gp60 sequence typing (Fig. 4, panel B). The gp60 sequence typing scheme presented with two genetically distinct branches. The major branch (most populous) showed extremely limited diversity, indicating they belong to the same gp60 subtype. Minor (least populous) branch showed substantially more diversity. Analysis using gp60 sequence typing showed extremely low diversity within the major branch, which comprised primarily of IIaA15G2R1 subtypes, with one IIaA16G3R1 subtype occupying its own minor branch (C393). The minor branch shows high diversity, with the remainder of the non-IIaA15G2R1 samples forming a distal group. Comparison between wgSNP and gp60 typing showed low correlation (*R*^*2*^=0.14).

**Figure 4.**
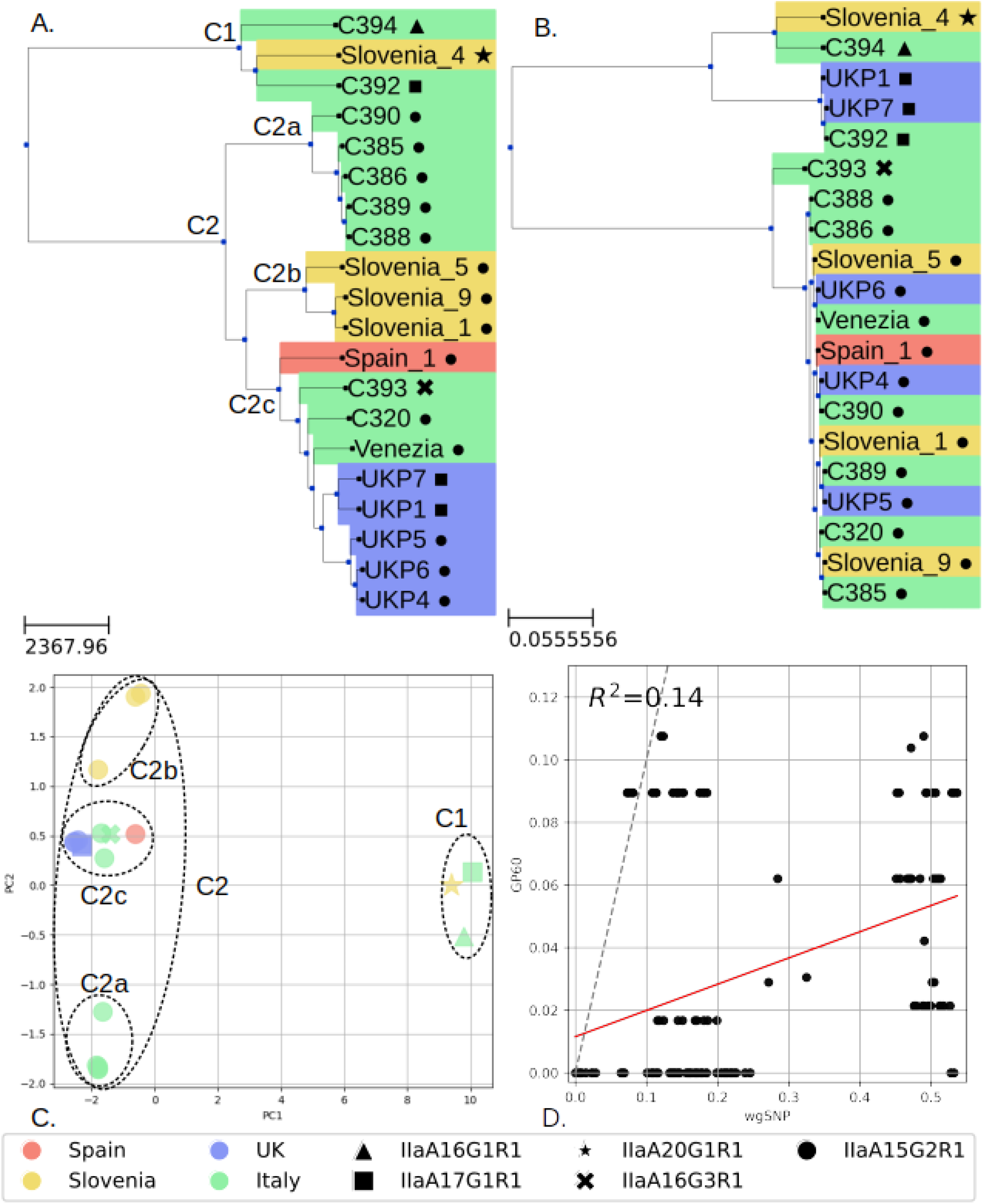
Phylogenetic analysis of this dataset using two typing schemes. (A) phylogenetic tree generated using the whole-genome SNP typing scheme, (B) phylogenetic tree generated using the gp60 sequences, (C) PCoA carried out using a Hamming distance matrix generated from the whole genome SNP set. Marker shapes represent gp60 subtype. Colourised by country of isolation. (D) Correlation analysis of typing schemes generated using the pairwise distance of each pair of samples within the dataset. The dashed line shows *x=y*, representing absolute parity between typing schemes. Cluster markers are placed by the root node in tree A for each cluster and used to annotate the PCoA plot.

## 4.0 Discussion

In the evaluation phase, the three clonal lineages (L1, L2 and L3) along with their sub-lineages were resolved by wgSNP Hamming distance analysis. Clustering by SNP distance (Fig. 2) indicated that sub-lineages within this dataset occupied a more distal location in SNP space. Interpretation of clustering results along with the results of MOI analysis (particularly using *Fws*) clearly resolved the clonal components of the simulated mixed samples. Heterogeneity plots can be used to give an indication of the number of populations which exist within a compound dataset, although the interpretation of such plots should be careful and taken in context with other data which may indicate the presence of multiple populations. PCoA plots of SNP distance can be used to reveal whether an ostensibly mixed sample could be a mixture of two known populations, such is the case with L1∪2, which showed strong signals of MOI (*Fws*>=1.0) and can be seen in Figure 2 to lie between lineages L1 and L2, reflecting the components of this mixture. The same conclusions can be made for L2∪3 (L2 and L3 mixture) and L1∪3 (L1 and L3 mixture). L1∪2∪3 (mixture of L1, L2 and L3) lay equidistant between Lineages 1, 2, and 3, demonstrating that Parapipe can be used resolve the individual components of a triple mixture. Results from running Parapipe on the real *C. parvum* IIa genomic dataset (see Table 1) showed substantial within-family diversity, revealed by using whole genome SNP analysis. MOI investigation of the dataset provided strong evidence of mixed infection (*Fws*>=0.95) in Venezia *(Fws*=1.0), and weak evidence of mixed infection (*Fws* 0.90-0.95) in C388 (*Fws*=0.922) and C389 (*Fws*=0.924). Corsi *et al*. reported a high multiplicity of infection (MOI) (Fws>=0.95) in both Venezia and C394 [26]. The apparent clonality of C349 (Fws=0.485) here is likely due to the fact that it contains the highest number of unique SNPs (814) in this dataset, making it the most genomically isolated sample. Since Fws represents the ratio of within-sample diversity to overall diversity across the dataset, samples from underrepresented populations or those with high genomic isolation tend to exhibit lower Fws values. Consequently, their polyclonality may be underestimated. It is therefore essential that interpretation of polyclonality using Fws be accompanied by phylogenetic analysis, and knowledge of sample origins. The limited metadata associated with these samples did not allow for further investigation into the incidence of MOI across different metadata categories, such as host or country. Phylogenetic analysis showed that there is accordance in the major clade structures between the two typing paradigms (Fig. 4), whereby two clades were defined. However, the membership of these clades differed substantially, potentially leading to inconsistent epidemiological interpretation. Group C1 in particular showed disparate results, whereby C394, C392 and Slovenia_4 all share a group under both typing schemes, but were grouped with UKP1 and UKP7 according to gp60, whereas wgSNP placed them with other UK isolates. The two typing schemes showed poor correlation within this dataset (R^2^=0.14), which is revealed by the pairwise correlation plot (Figure 4D). Furthermore, almost all points in the correlation analysis land below the *x=y* line, indicating a lower level of typing resolution achievable by using gp60. Samples which were closely associated by wgSNP (such as UK isolates) were split into two distinct gp60 subtypes, IIaA15G2R1 and IIaA17G1R1, indicating a discrepancy between local and global genome mutation and recombination rates. Geographical origins were more robustly recovered by wgSNP than gp60 typing. This is likely due to the reduced resolution of a single locus Sanger sequence typing approach, which relies on variation embedded in a very small subset of the *Cryptosporidium* genome. Mutations relevant to elucidating transmission, geographic origin, and phenotype which are embedded within regions not covered by a molecular typing scheme will not influence the topology of the trees they are used to generate but will be captured by a wgSNP typing approach. Furthermore, a greater degree of discrimination was achieved using wgSNP, resulting in deeper leaf nodes within the tree. Typing by gp60 places two UK isolates, UKP7 and UKP1, with the Italian isolate C392, since they belong to the same subtype (IIaA17G1R1). However, analysis by wgSNP placed all UK isolates together, and showed considerable distance between all UK isolates and C392, indicating substantial genomic divergence between these isolates which is not captured by gp60 typing.

**Table 1.** Mapping, MOI, and SNP calling results from running the *C. parvum* gp60 *family* IIa dataset through Parapipe. DOC = Depth of Coverage; BOC>=5x = The percentage of the reference genome covered to a minimum depth of 5x; Norm. GG area = the normalised area under the Gini-Granularity curve [25]; *Fws* = the within sample F statistic; SNPs: total (unique) = the total and unique number of SNPs detected in this sample.

| ID | Mean DOC | BOC≥5x | Norm GG area | Fws | SNPs: total (unique) |
| --- | --- | --- | --- | --- | --- |
| **C320** | 674.6 | 99.93 | 0.993 | 0.744 | 1120 (99) |
| **C385** | 619.2 | 99.95 | 0.992 | 0.828 | 1376 (4) |
| **C386** | 555.2 | 99.95 | 0.992 | 0.865 | 1392 (6) |
| **C388** | 606.7 | 99.96 | 0.993 | 0.922 | 1397 (0) |
| **C389** | 640.9 | 99.95 | 0.993 | 0.924 | 1403 (3) |
| **C390** | 379.4 | 99.97 | 0.991 | 0.582 | 1327 (41) |
| **C392** | 317.8 | 99.97 | 0.991 | 0.491 | 4264 (524) |
| **C393** | 450.9 | 99.97 | 0.992 | 0.409 | 1182 (90) |
| **C394** | 386.1 | 99.94 | 0.991 | 0.485 | 4188 (814) |
| **Venezia** | 973.1 | 99.89 | 0.991 | 1.003 | 1147 (100) |
| **Slovenia_1** | 476.2 | 99.96 | 0.992 | 0.815 | 1603 (24) |
| **Slovenia_4** | 286.3 | 99.95 | 0.991 | 0.54 | 4025 (681) |
| **Slovenia_5** | 351.1 | 99.96 | 0.991 | 0.857 | 1155 (34) |
| **Slovenia_9** | 552.8 | 99.94 | 0.991 | 0.813 | 1644 (24) |
| **Spain_1** | 564.4 | 99.92 | 0.992 | 0.791 | 1314 (449) |
| **UKP1** | 875.7 | 99.92 | 0.984 | 0.732 | 754 (79) |
| **UKP4** | 260.2 | 99.37 | 0.948 | 0.874 | 860 (29) |
| **UKP5** | 37.1 | 98.96 | 0.945 | 0.782 | 857 (65) |
| **UKP6** | 147.4 | 99.97 | 0.966 | 0.881 | 871 (21) |
| **UKP7** | 104.6 | 96.96 | 0.949 | 0.758 | 750 (89) |

These results indicated that gp60 typing can resolve major structures within *Cryptosporidium* population data. However, it was less sensitive than a wgSNP approach at resolving highly granular population structure, particularly across closely related samples. wgSNP analysis indicated more within-clade diversity than the gp60 typing scheme due to higher resolution. This furnishes the epidemiologist with higher resolution data with which to draw epidemiological inferences and elucidate transmission dynamics, which may be lost with lower resolution typing schemes.

A comparison between functionality of Parapipe and similar publicly available pipelines (see Table 2) highlighted that no other pipeline carries out the full repertoire of pre-assembly analysis that Parapipe is capable of. MOI analysis is an area of analysis missing from other pipelines within the cohort, which were historically developed to analyse subsamples from clonal pathogen populations. Furthermore, the generation of a binary allelic matrix by Parapipe facilitates an enormous amount of downstream analysis, such as population structure analysis, phylogenetic tree reconstruction, genetic diversity assessment, association studies to link genetic variants with phenotypes, detection of selective sweeps, admixture analysis, identification of conserved or unique genomic regions across populations, and machine learning applications for genotype-phenotype prediction or classification.

**Table 2.**
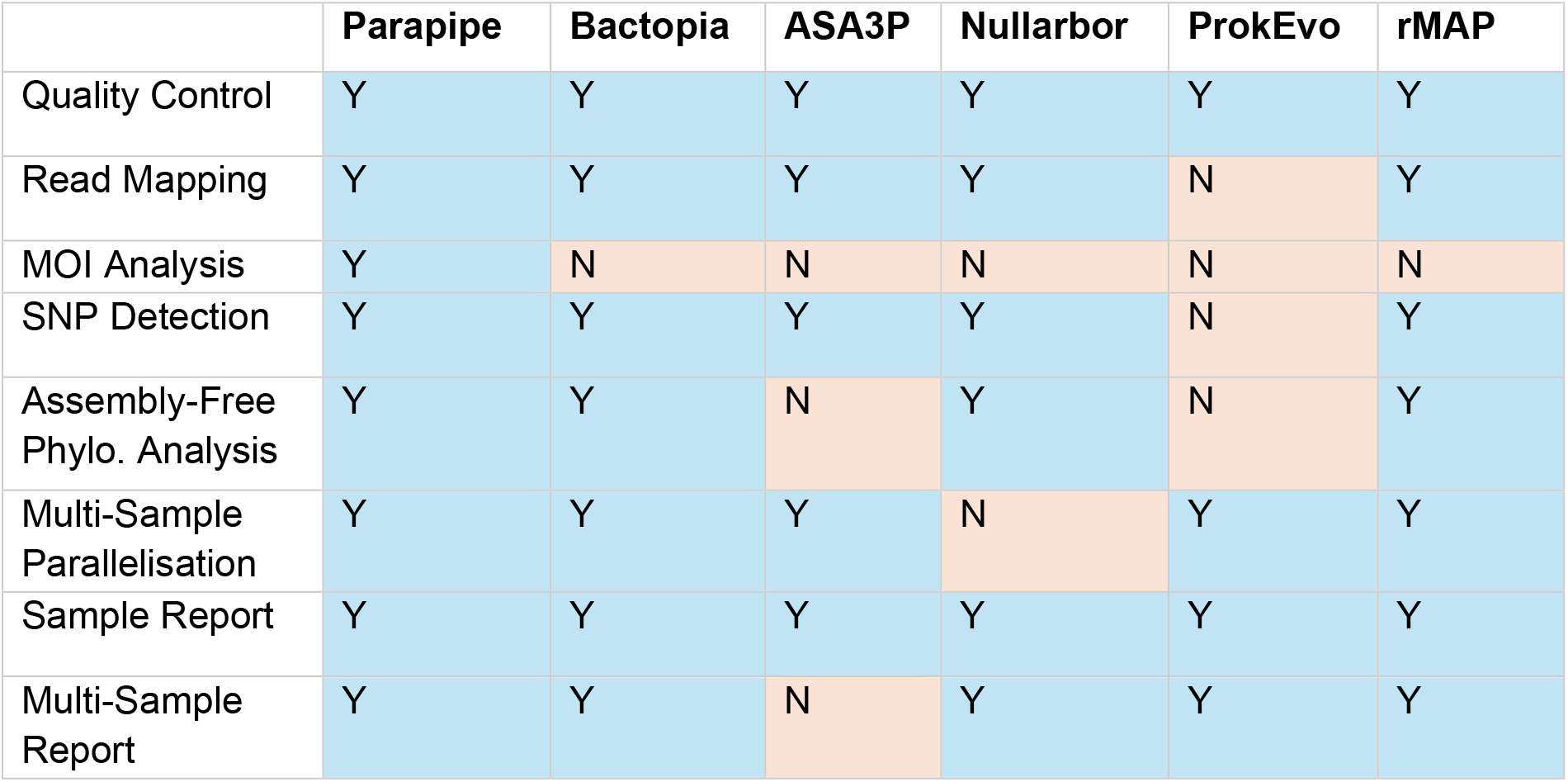
A comparison between publicly available pipelines for processing pathogen NGS data and carrying out assembly-free analysis.

|  | Parapipe | Bactopia | ASA3P | Nullarbor | ProkEvo | rMAP |
| --- | --- | --- | --- | --- | --- | --- |
| Quality Control | Y | Y | Y | Y | Y | Y |
| Read Mapping | Y | Y | Y | Y | N | Y |
| MOI Analysis | Y | N | N | N | N | N |
| SNP Detection | Y | Y | Y | Y | N | Y |
| Assembly-Free Phylo. Analysis | Y | Y | N | Y | N | Y |
| Multi-Sample Parallelisation | Y | Y | Y | N | Y | Y |
| Sample Report | Y | Y | Y | Y | Y | Y |
| Multi-Sample Report | Y | Y | N | Y | Y | Y |

Currently, due to the expensive and time-consuming nature of carrying out NGS on all clinical samples of *Cryptosporidium*, along with the demonstrable utility of a MLVA scheme for identifying and investigating outbreaks [8], [9], it is likely that the application of NGS and wgSNP analysis will be limited to selected clinical samples (e.g. those taken from outbreak events) until a scalable and cost-effective laboratory workflow has been designed and implemented. Despite this limitation, it is likely that the future of *Cryptosporidium* typing will utilise NGS technologies, enabled by improvements in sample and library preparation, allowing analyses based on the wgSNPs distance rather than interrogation of a small subset of VNTR loci. This developmental leap has already been made for other pathogens, such as *Salmonella enterica* and *Mycobacterium tuberculosis* [27], [28]. Parapipe generates an allelic presence/absence matrix from all samples within the dataset, along with all the data required to perform a thorough quality control assessment for each sample prior to their incorporation into a database of historical samples. Such a database can then be queried to yield key phylogenomic information, informing about the relationship between *Cryptosporidium* samples based on their exhibited allelic repertoire. NGS enables high-resolution and robust analysis of MOI, a phenomenon with potentially significant epidemiological implications for *Cryptosporidium*. MOI analysis can identify mixed infections, potentially signalling exposure to multiple or overlapping transmission sources, differentiate between local transmission and importation, and assess its role in the emergence of new genotypes. While the dataset used in this study lacks the epidemiological metadata required to elucidate specific transmission pathways, the MOI findings underscore the complexity of *Cryptosporidium* infections.

Parapipe was written to sit in the middle of a larger workflow which can be employed by public health laboratories, where the first step is *in silico* species identification to facilitate reference genome selection, and downstream analysis includes exposing output to a database, and epidemiological hypothesis testing. Parapipe streamlines genomic analysis, providing public health laboratories with a tool to integrate MOI data with geographic, demographic, and environmental variables. This integration has the potential to enhance the epidemiological precision of outbreak investigations, facilitating an effective public health response.

## 5.0 Conclusion

In this paper, we have presented Parapipe, a novel bioinformatics pipeline designed to be the foundation of a more complex analytical framework for pathogenic protozoan NGS data. The generation of an allele matrix, along with allele-frequency and MOI data, facilitates in depth phylogenetic analysis. Parapipe streamlines and standardises a bespoke bioinformatic workflow designed to carry out quality control, heterogeneity, and phylogenetic analysis of protozoan NGS data, which are essential parts of any characterisation process intended to enable tracking and control of disease in humans or animals. Here, we use both complex simulated datasets and a case study using real data to show that Parapipe is capable of automating quality control and reporting, identifying mixtures within NGS datasets, and carrying out both MOI investigation and whole genome SNP typing for extracting crucial information necessary for determining the phylogenetic relationship between samples. Parapipe represents, to our knowledge, the first publicly available pipeline designed for the analysis and handling of such data, and benefits from development within a public health environment, building a modular pipeline that is ISO accreditable by design. Parapipe has been thoroughly tested and validated, using both individual module and end-to-end testing approaches. It is entirely modularised using Nextflow with dependencies managed through the use of Singularity containers. The modularity of the pipeline facilitates the implementation of new functionalities into Parapipe. It is a crucial first step in the development of a full suite of robust, validated bioinformatic tools, which can be used to aid the public health response to *Cryptosporidium*. It represents the first contact point for dealing with clinical (human or animal) or environmental NGS data, giving a detailed but easily digestible overview of the data. Parapipe was designed in accordance with the needs of public health agencies and laboratories.

## Data Availability

The source code for Parapipe is available at https://github.com/ArthurVM/Parapipe. The clonal and heterogeneous lineage definition files along with scripts used to generate lineage NGS datasets are available at https://github.com/ArthurVM/Parapipe_test_data. The source code for pyMOI is available at https://github.com/ArthurVM/PyMOI. Scripts for simulating NGS reads using ART can be found at https://github.com/ArthurVM/ngsContrive.

## Acknowledgements

We are grateful to Martin Swain, Deborah Oladele, and the members of the Pathogen Genomics Unit for their feedback in the use of Parapipe during the testing and development phase.

## Author Contributions Statement

AM conceived the project with RC, GR, and SC. AM designed, developed, and tested Parapipe, and wrote the manuscript. TC aided with design and advised on practical implementation within a public health context. All authors contributed to the broad design of Parapipe, manuscript proof reading and editing.

