## supplementary materials for "Parapipe: a Pipeline for Handling Parasite NGS Datasets and its Application to Cryptosporidium"

Testing Modules were designated from the processes detailed in Fig. 1. Six discrete modules were defined, which are detailed in Table S1.

| Module Code | Processes | Function | Dataset Code | Description | Intended Result | Actual Result |
|---|---|---|---|---|---|---|
| TM01 | 1.2 | Ensure the process which validates the input data functions as intended. | TM01.1 | Empty FQ | FAIL | FAIL |
| | | | TM01.2 | Malformed FQ | FAIL | FAIL |
| TM02 | 1.3 | Ensure the process which ensures there are enough reads in the input dataset functions as intended. | TM02.1 | 10% ref 10x cov | FAIL | FAIL |
| | | | TM02.2 | 499,999 read pairs | FAIL | FAIL |
| | | | TM02.3 | 500,000 read pairs | PASS | PASS |
| | | | TM02.4 | 500,001 read pairs | PASS | PASS |
| TM03 | 1.4 | Ensure the process which filters out short, low quality, and low complexity reads functions as intended. | TM03.1 | 499,999 read pairs with 2 50n read pairs designed to be filtered out | FAIL | FAIL |
| | | | TM03.2 | 499,999 read pairs with 2 low quality read pairs designed to be filtered out | FAIL | FAIL |
| | | | TM03.3 | 499,999 read pairs with 2 low complexity (poly A) read pairs designed to be filtered out | FAIL | FAIL |
| TM04 | 1.7 | Ensure mapping statistics are being reported correctly. | TM04.1 | 1% ref at 25x | Correct mapping statistics | Median DOC: 25.0 Ref cov: 0.00998 |
| | | | TM04.2 | 10% ref at 25x | Correct mapping statistics | Median DOC: 25.0 Ref cov: 0.09998 |
| | | | TM04.3 | 25% ref at 25x | Correct mapping statistics | Median DOC: 25.0 Ref cov: 0.24998 |
| | | | TM04.4 | 50% ref at 25x | Correct mapping statistics | Median DOC: 25.0 Ref cov: 0.50063 |
| | | | TM04.5 | 75% ref at 25x | Correct mapping statistics | Median DOC: 25.0 Ref cov: 0.75063 |
| | | | TM04.6 | 100% ref at 25x | Correct mapping statistics | Median DOC: 25.0 Ref cov: 0.99998 |
| | | | TM04.7 | 100% ref at 1x | Correct mapping statistics | Median DOC: 2..0 Ref cov: 0.48645 |
| TM05 | 1.8-2.1 | Ensure heterozygosity is being correctly and robustly reported. | TM05.1 | 2 populations introduced across 100 sites. AF={0.9, 0.1} | 2 clusters centered at correct AF. Fws >= 0.95 | 2 clusters with correct AF. |
| | | | TM05.2 | 2 populations introduced across 1000 sites. AF={0.9, 0.1} | 2 clusters centered at correct AF. Fws >= 0.95 | 2 clusters with correct AF |
| | | | TM05.3 | 3 populations introduced across 100 sites. AF={0.9, 0.6, 0.1} | 3 clusters centered at correct AF. Fws >= 0.95 | 3 clusters with correct AF |
| | | | TM05.4 | 3 populations introduced across 1000 sites. AF={0.9, 0.6, 0.1} | 3 clusters centered at correct AF. Fws >= 0.95 | 3 clusters with correct AF |
| TM06 | 2.2-2.3 | Ensure samples are being correctly characterised according to their phylogenetic position. | TM06.1 | 10 datasets with defined hierarchical lineage structure down to 2nd order | 3 clusters each containing 4 samples. Clustered by Lineage, with 1 distal branch to each cluster | Correct tree and cluster topology. |

Table S1. Testing modules and datasets used to test Parapipe. AF=allele frequency, as a ratio of reference to alternative allele at a given site. Results of Module testing are included in the Actual Result column.

| Order 1 Lineage | Description | Order 2 Lineage | Description |
|---|---|---|---|
| L1 | 5 SNPs on each chromosome. | L1.1 | 5 further SNPs on each chromosome. |
| | | L1.2 | |
| | | L1.3 | |
| | | L1.4 | |
| | | L1.5 | |
| L2 | 5 SNPs on each chromosome. | L2.1 | 5 further SNPs on each chromosome. |
| | | L2.2 | |
| | | L2.3 | |
| | | L2.4 | |
| | | L2.5 | |
| L3 | 5 SNPs on each chromosome. | L3.1 | 5 further SNPs on each chromosome. |
| | | L3.2 | |
| | | L3.3 | |
| | | L3.4 | |
| | | L3.5 | |

Table S2. Simulated datasets used to test module TM06.

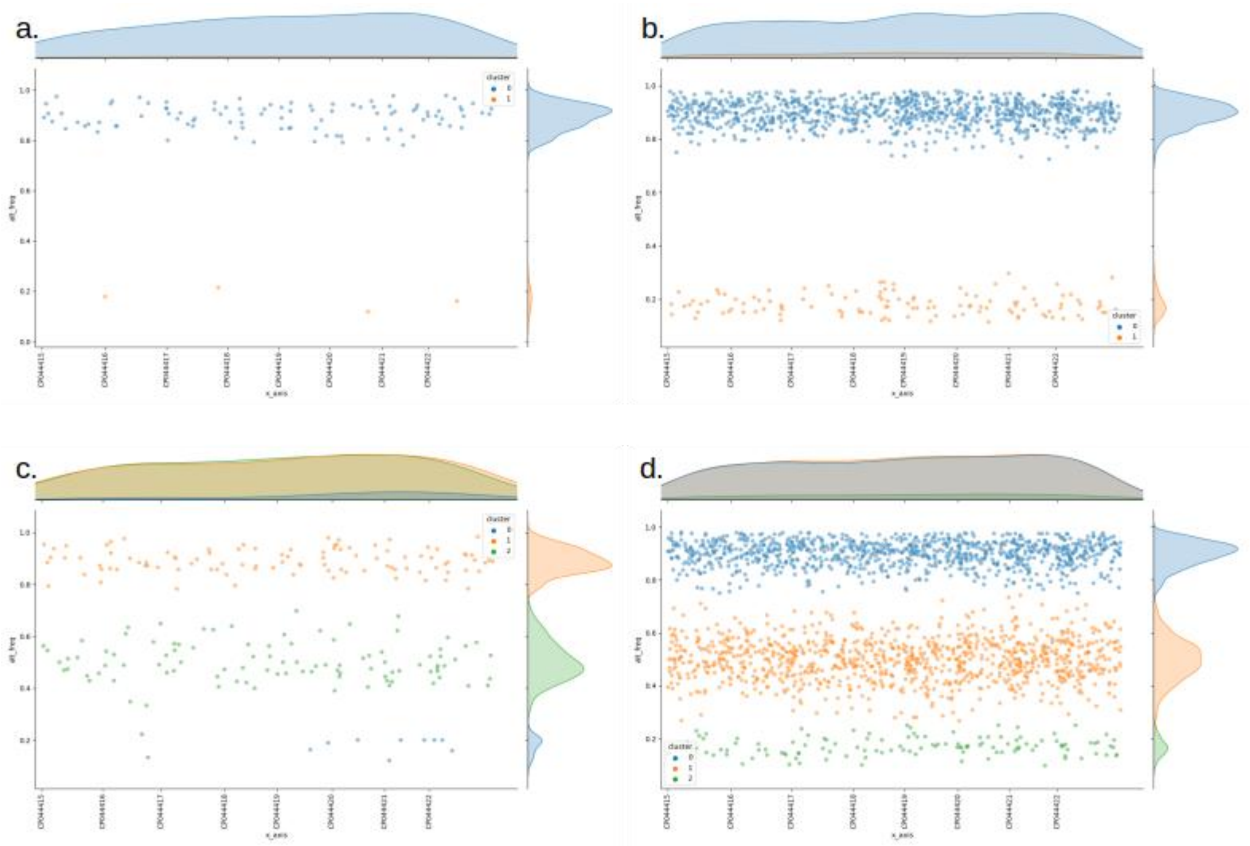


Figure S1. MOI plots generated from running testing module TM05 using the dataset TM05.1 - TM05.4.

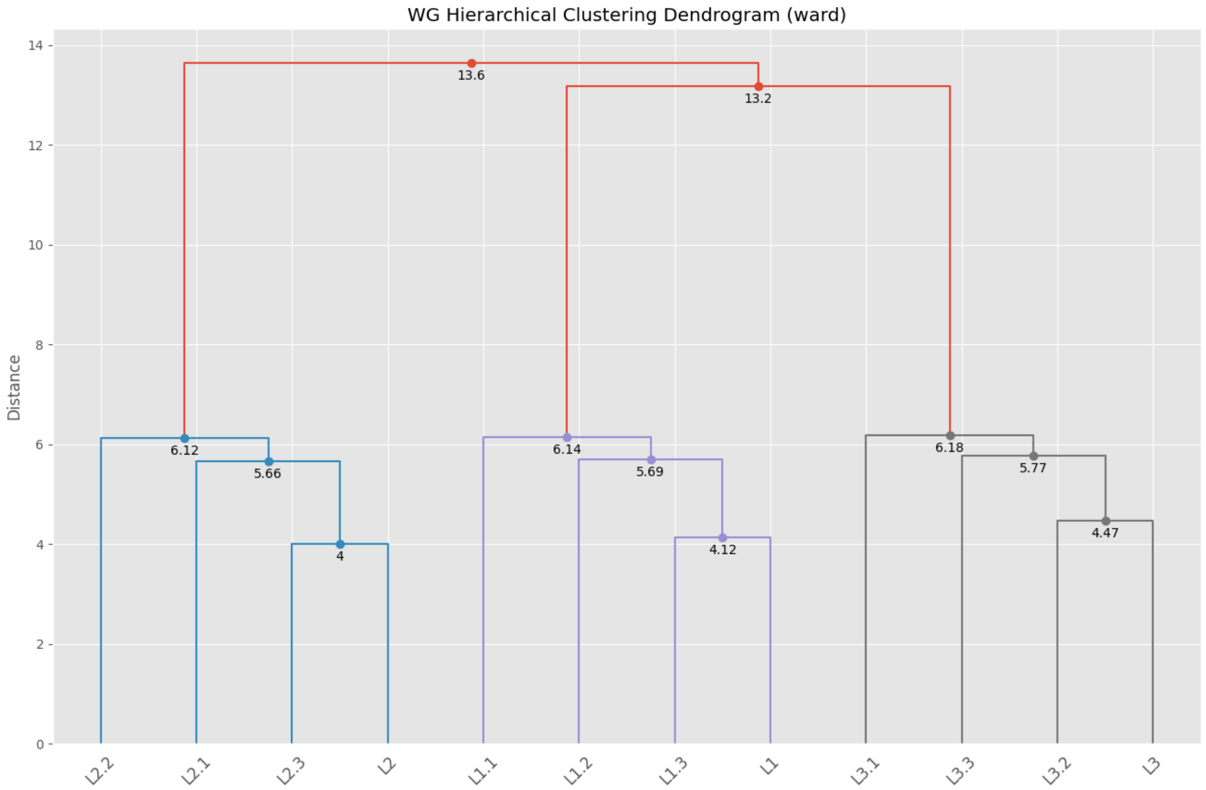

Figure S2. A wgSNP tree generated by running testing module TM06 on test dataset TM06.1.

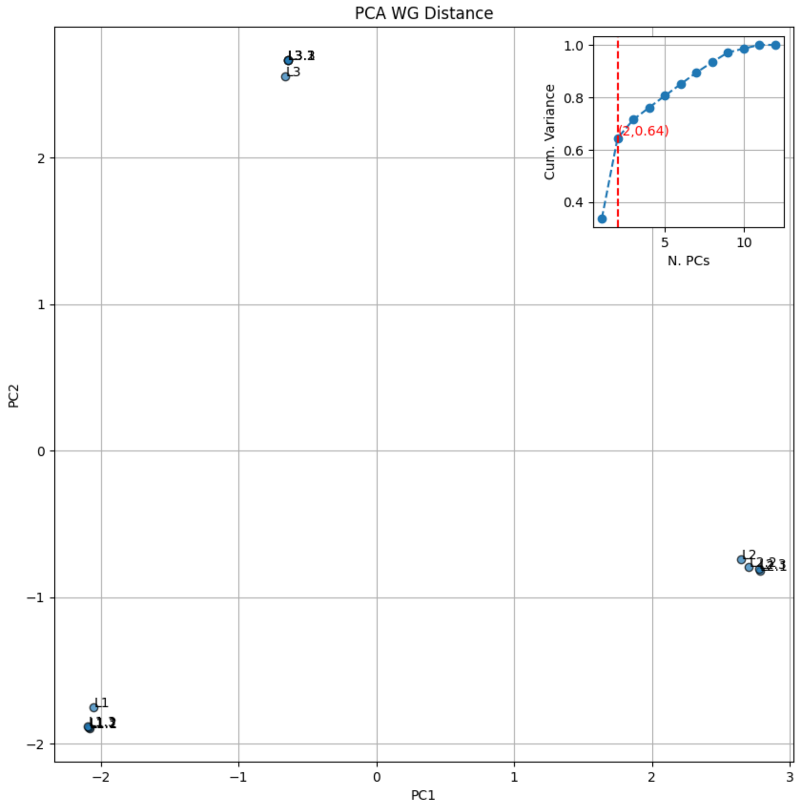


Figure S3. A PCoA plot of wgSNP distance generated by running testing module TM06 on test dataset TM06.1

| ID | Mean DOC | Median DOC | BOC>= 5x | GG area | Norm GG area | Av. Qual | Av. Insert Size | Fws | SNPs: total (unique) |
|---|---|---|---|---|---|---|---|---|---|
| **L3.1** | 19.8 | 20 | 99.93 | 0.102 | 0.935 | 36.4 | 199.3 | 0 | 78 (42) |
| **L2.4** | 19.8 | 20 | 99.92 | 0.103 | 0.935 | 36.4 | 199.3 | 0.002 | 81 (40) |
| **L1.4** | 19.8 | 20 | 99.92 | 0.101 | 0.935 | 36.4 | 199.3 | 0 | 74 (37) |
| **L3** | 19.8 | 20 | 99.93 | 0.102 | 0.935 | 36.4 | 199.3 | 0.003 | 36 (0) |
| **L1.1** | 19.8 | 20 | 99.92 | 0.102 | 0.935 | 36.4 | 199.3 | 0 | 76 (38) |
| **L3.5** | 19.8 | 20 | 99.91 | 0.102 | 0.935 | 36.4 | 199.3 | 0 | 74 (38) |
| **L1** | 19.8 | 20 | 99.93 | 0.102 | 0.935 | 36.4 | 199.3 | 0 | 38 (0) |
| **L3.3** | 19.8 | 20 | 99.92 | 0.102 | 0.935 | 36.4 | 199.3 | 0.004 | 72 (36) |
| **L2.2** | 19.8 | 20 | 99.92 | 0.102 | 0.935 | 36.4 | 199.3 | 0 | 80 (38) |
| **L2.5** | 19.8 | 20 | 99.93 | 0.102 | 0.935 | 36.4 | 199.3 | 0.002 | 80 (38) |
| **HL1-2** | 39.7 | 39 | 99.99 | 0.072 | 0.953 | 36.4 | 199.3 | 1.119 | 80 (0) |
| **L2.1** | 19.9 | 20 | 99.92 | 0.102 | 0.935 | 36.4 | 199.3 | 0 | 80 (38) |
| **L2** | 19.8 | 20 | 99.92 | 0.102 | 0.935 | 36.4 | 199.3 | 0 | 42 (0) |
| **L2.3** | 19.9 | 20 | 99.91 | 0.102 | 0.935 | 36.4 | 199.3 | 0 | 78 (36) |
| **L3.2** | 19.8 | 20 | 99.93 | 0.101 | 0.935 | 36.4 | 199.3 | 0 | 74 (38) |
| **HL2-3** | 39.7 | 39 | 99.99 | 0.072 | 0.952 | 36.4 | 199.3 | 1.087 | 78 (0) |
| **L1.3** | 19.8 | 20 | 99.91 | 0.101 | 0.935 | 36.4 | 199.3 | 0 | 75 (37) |
| **L1.2** | 19.8 | 20 | 99.92 | 0.102 | 0.935 | 36.4 | 199.3 | 0 | 75 (37) |
| **HL1-3** | 39.7 | 39 | 99.99 | 0.072 | 0.952 | 36.4 | 199.3 | 1.107 | 74 (0) |
| **L1.5** | 19.9 | 20 | 99.91 | 0.102 | 0.935 | 36.4 | 199.3 | 0 | 69 (31) |
| **HL1-2-3** | 59.5 | 59 | 100 | 0.059 | 0.96 | 36.4 | 199.3 | 1.008 | 115 (0) |
| **L3.4** | 19.8 | 20 | 99.93 | 0.101 | 0.935 | 36.4 | 199.3 | 0 | 74 (38) |

Figure S5. The table of mapping statistics for all samples presented in the collated run report.

```
.
└── My_Run_Directory/
    ├── bams/
    │   ├── sample1_grouped.bam
    │   ├── sample2_grouped.bam
    │   ├── sample3_grouped.bam
    │   └── ...
    ├── sample1/
    │   ├── sample1.err
    │   ├── sample1_report.json
    │   ├── sample1_report.pdf
    │   ├── MOI/
    │   │   └── ... heterozygosity data
    │   ├── preprocessing/
    │   │   └── ... mapping and QC data
    │   ├── raw_read_QC_reports/
    │   │   └── sample1_fastp.json
    │   └── VariantAnalysis/
    │       └── SNP/
    │           └── sample1.vcf.gz
    ├── sample2/
    │   └── ...
    ├── sample3/
    │   └── ...
    ├── ...
    ├── mapping_stats/
    │   ├── sample1_mapstats.json
    │   ├── sample2_mapstats.json
    │   ├── sample3_mapstats.json
    │   └── ...
    ├── multiqc_report.html
    ├── Parapipe_report.html
    ├── Parapipe_report.pdf
    ├── phylo/
    │   ├── allele_matrix.csv
    │   ├── sample1.snps.bed
    │   ├── sample2.snps.bed
    │   ├── sample3.snps.bed
    │   └── ...
    └── REFDATA/
        ├── ref.fasta
        └── ref.gff
```

Figure S7. The directory structure for a Parapipe run output directory. The run report is captured in the Parapipe_report.html file, and sample report pdfs deposited in their sample directories.

| ID | gp60 Subtype | Country | BioProject | Accession | Study |
|---|---|---|---|---|---|
| **C393** | IIaA16G3R1 | Italy | PRJNA633764 | SRR11817809 | Corsi et al. (2023) |
| **C392** | IIaA17G1R1 | Italy | PRJNA633764 | SRR11817810 | Corsi et al. (2023) |
| **C390** | IIaA15G2R1 | Italy | PRJNA633764 | SRR11817812 | Corsi et al. (2023) |
| **C389** | IIaA15G2R1 | Italy | PRJNA633764 | SRR11817813 | Corsi et al. (2023) |
| **C388** | IIaA15G2R1 | Italy | PRJNA633764 | SRR11817814 | Corsi et al. (2023) |
| **C386** | IIaA15G2R1 | Italy | PRJNA633764 | SRR11817815 | Corsi et al. (2023) |
| **C385** | IIaA15G2R1 | Italy | PRJNA633764 | SRR11817816 | Corsi et al. (2023) |
| **Venezia** | IIaA15G2R1 | Italy | PRJNA633764 | SRR11817817 | Corsi et al. (2023) |
| **C394** | IIaA16G1R1 | Italy | PRJNA633764 | SRR11817821 | Corsi et al. (2023) |
| **C320** | IIaA15G2R1 | Italy | PRJNA633764 | SRR11817823 | Corsi et al. (2023) |
| **Spain_1** | IIaA15G2R1 | Spain | PRJNA634014 | SRR11818073 | Corsi et al. (2023) |
| **Slovenia_9** | IIaA15G2R1 | Slovenia | PRJNA634014 | SRR11818074 | Corsi et al. (2023) |
| **Slovenia_5** | IIaA15G2R1 | Slovenia | PRJNA634014 | SRR11818076 | Corsi et al. (2023) |
| **Slovenia_4** | IIaA20G1R1 | Slovenia | PRJNA634014 | SRR11818077 | Corsi et al. (2023) |
| **Slovenia_1** | IIaA15G2R1 | Slovenia | PRJNA634014 | SRR11818079 | Corsi et al. (2023) |
| **UKP4** | IIaA15G2R1 | UK | PRJNA253843 | SRR6147581 | Hadfield et al. (2015) |
| **UKP5** | IIaA15G2R1 | UK | PRJNA253843 | SRR6147587 | Hadfield et al. (2015) |
| **UKP6** | IIaA15G2R1 | UK | PRJNA253843 | SRR6147945 | Hadfield et al. (2015) |
| **UKP7** | IIaA17G1R1 | UK | PRJNA253843 | SRR6147964 | Hadfield et al. (2015) |
| **UKP1** | IIaA17G1R1 | UK | PRJNA253843 | SRR6871415 | Hadfield et al. (2015) |

Figure S8. The dataset consisting of *C. parvum* samples belonging to gp60 subtype family IIa.
